# *VistoSeg*: processing utilities for high-resolution Visium/Visium-IF images for spatial transcriptomics data

**DOI:** 10.1101/2021.08.04.452489

**Authors:** Madhavi Tippani, Heena R. Divecha, Joseph L. Catallini, Sang Ho Kwon, Lukas M. Weber, Abby Spangler, Andrew E. Jaffe, Stephanie C. Hicks, Keri Martinowich, Leonardo Collado-Torres, Stephanie C. Page, Kristen R. Maynard

## Abstract

**Background:** Spatial transcriptomics is a next-generation sequencing technology that combines the strengths of transcriptome-wide RNA-sequencing with histological imaging to generate spatial maps of gene expression in intact tissue sections. The 10x Genomics Visium and Visium-Immunofluorescence (Visium-IF) platforms are widely available commercial technologies for quantifying spatially-resolved gene expression. These technologies directly couple gene expression with high resolution histological or immunofluorescence images that contain rich morphological information about the tissue section. However, extracting and integrating image features with gene expression data remains challenging.

**Results:** Using MATLAB, we developed *VistoSeg*, which is a pipeline to process, analyze, and interactively visualize the high-resolution images from the 10x Genomics Visium and Visium-IF platforms. The output from *VistoSeg* can then be integrated with the spatial-molecular information in downstream analyses using common programming languages, such as R or Python.

**Conclusion:** *VistoSeg* provides user-friendly tools for integrating image-derived metrics from histological and immunofluorescent images with spatially-resolved gene expression data. This integrated approach can advance our understanding of the transcriptional landscape within tissue architecture. *VistoSeg* is freely available at http://research.libd.org/VistoSeg/.

**Impact Statement:** Technologies for measuring gene activity levels, referred to as gene expression, have been evolving over decades and are the core of the transcriptomics subfield within genomics. The first report describing individual cell gene expression is from 2009 and as a method it became commercially available in 2014. While single cell transcriptomics increased our resolution beyond homogenate tissue, the advent of spatial transcriptomics technologies and commercial availability of spatial gene expression platforms, such as Visium, has facilitated studying gene expression in anatomical context. Visium measures local gene expression within the histological organization of single 6.5 mm^2^ cryosection of tissue. Spatially-resolved transcriptomics provides a new challenge: integrating spatial gene expression with high resolution tissue images (brightfield histology or fluorescent antibody staining). *VistoSeg* image processing software is compatible with both Visium and Visium-IF from 10x Genomics, which are spatially-resolved transcriptomics assays employing histological and immunofluorescent images, respectively. From these images, the number of cells, identity of cell types, and other image-derived markers can be obtained for thousands of 2,375 µm^2^ spots, where genome-wide gene expression is also measured. *VistoSeg* provides tools that enable processing these images in the context of gene expression maps to integrate these two high dimensional data types, and thus help unlock the new frontier in transcriptomics.

## Introduction

In the last decade, RNA sequencing (RNA-seq) has moved beyond whole-tissue resolution to single cells or nuclei RNA-seq. While this leap forward motivated new methods and biological questions, the spatial arrangement of individual cell types within the tissue remained underexplored. Spatially-resolved transcriptomics (1) is a new class of technologies that measures gene expression along two-dimensional spatial coordinates. 10x Genomics provided the first widely used commercially-available spatial transcriptomics platform, Visium, for generating transcriptome-wide spatial gene expression data in intact tissue sections. Because spatial transcriptomics data includes paired gene expression and microscopy images from the same tissue slice, a major goal of the field has been to integrate data from these complementary modalities.

The 10x Genomics Visium spatial gene expression workflow uses an on-slide spatial barcoding strategy to map RNA-seq reads to defined anatomical locations (‘spots’) in an intact tissue section (**Fig. 1A**). In Visium, each slide contains four arrays (**Fig. 1B**: A1, B1, C1, D1) with each array containing ∼5,000 gene expression capture spots that are 55µm in diameter (2,375 µm^2^ in area) and spaced 100µm center-to-center in a honeycomb pattern (**Fig. 1B**). Using cDNA synthesis on the Visium assay, coupled with RNA-seq, gene expression measurements for these spots are obtained (**Fig. 1C**). The platform currently supports two major workflows: Visium and Visium-Immunofluorescence (IF), which is referred to as Visium-IF hereinafter. In the Visium workflow, tissue sections are stained with hematoxylin and eosin (H&E), and a brightfield histological image is acquired. In Visium-IF, tissue sections are labeled with antibodies conjugated to fluorophores, and multiplex fluorescent images are acquired to visualize proteins of interest. These images are used to create an integrated map of transcriptome-wide gene expression within the tissue architecture. In the case of Visium-IF, transcriptomic data can also be analyzed in the context of proteomic data from antibody labeling.

**Figure 1:**
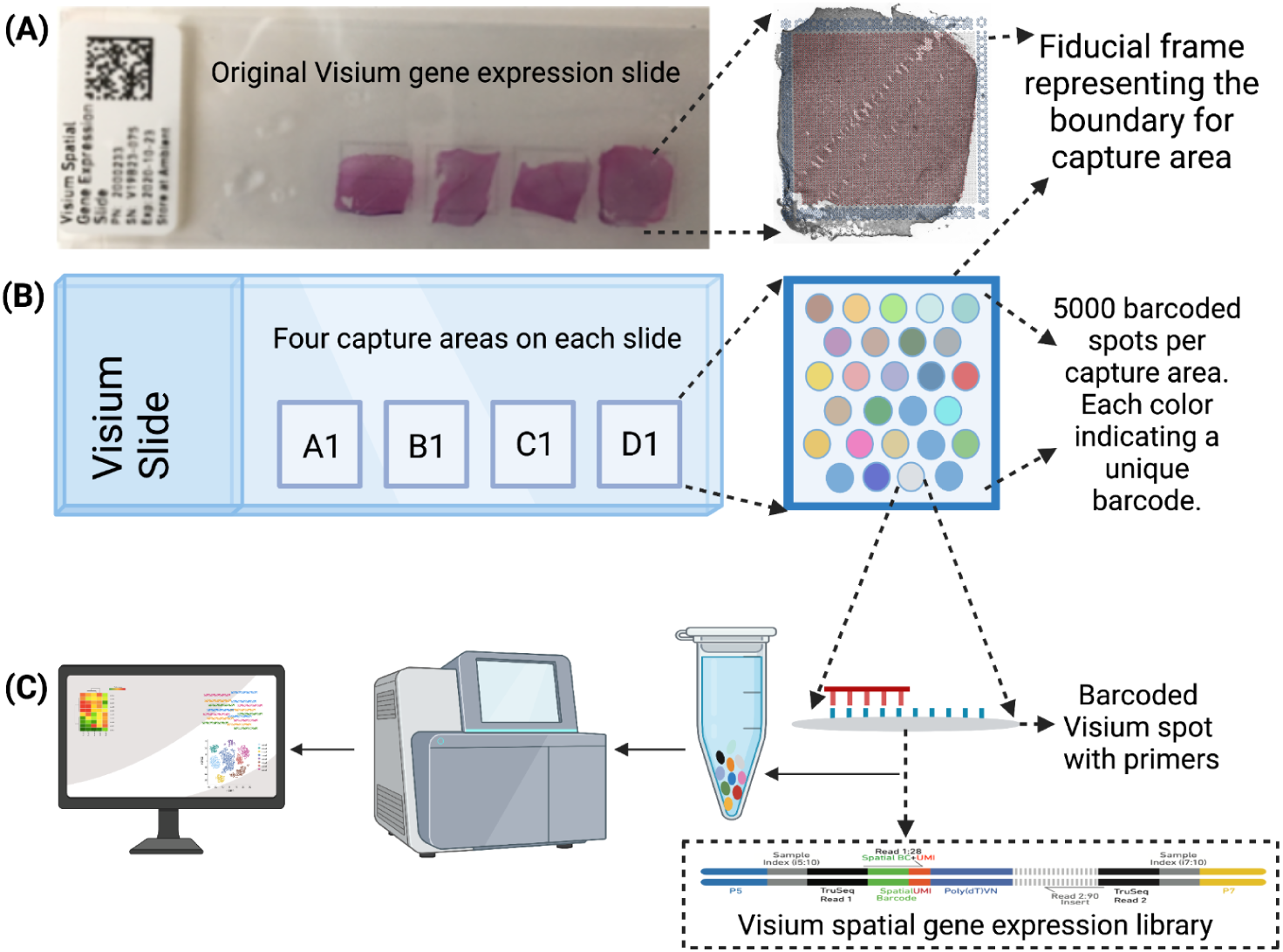
Overview. (**A**) Original ‘Visium spatial gene expression slide’ containing 4 arrays. Each array contains a 6.5 mm x 6.5 mm capture area, which is bounded by a fiducial frame. (**B**) Each capture area contains a grid with ∼5000 unique spatially barcoded spots with capture probes to collect the mRNA information from the tissue section collected on the slide. (**C**) Barcodes indicating the location of each spot are incorporated during on-slide cDNA synthesis. The cDNA is then removed from the slide, prepared into libraries, and sequenced. This allows each read to be mapped back to the correct spatial coordinates on the histological image, providing a transcriptome-wide readout of gene expression at each spatial coordinate.

To maintain RNA integrity for compatibility with the downstream transcriptomic workflow, speed and throughput are important considerations for Visium image acquisition. Thus, slide scanning microscopy is often utilized for Visium imaging; however, these microscopes routinely export imaging data as a single large file, necessitating software to split large whole slide images into individual arrays. Moreover, commercial image analysis software for Visium (2) or pathology imaging software (3) do not support quantitative analysis of Visium images for integration with molecular gene expression data. However, these images represent a rich, untapped resource for downstream analyses. For example, nuclei segmentation of H&E images can be used to estimate the number of nuclei in each Visium spot or to identify cell types based on classic morphologies (4,5). Analysis of histological images can also identify spots containing a single cell, or spots enriched in specific cellular compartments (i.e. axon and dendrite-rich neuropil in brain tissue) (6).

On the other hand, Visium-IF (7), is a high-throughput multi-omics approach that combines proteomics and transcriptomics analysis to facilitate the correlation of protein abundance measurements to gene expression. The workflow for Visium-IF is identical to Visium, with the exception of upstream tissue staining and imaging. Instead of H&E staining used in Visium, Visium-IF uses fluorescent antibody staining to label proteins of interest. Tissue sections are then fluorescently imaged with each protein assigned a unique color and distinct channel in a multiplex image. Analysis of multiplex fluorescent images can provide detailed proteomic maps that can be used to quantify important cellular or pathological features for integration with transcriptomic data (8). For example, images can be used to quantify pathological hallmarks of various diseases, which can then be linked with gene expression data to better understand molecular associations with disease pathology (9). Using known proteomic cell type markers, cell types can be labeled for identifying the cell populations associated with gene expression in a given Visium spot. *In silico* methods that aim to identify cell type proportions at the spot level are called ‘spot deconvolution’ algorithms and are typically based either on spatially-resolved gene expression data only or also use the cell morphology derived from H&E images (10–12). Visium-IF can thus help generate a ground truth for evaluating cell type spot deconvolution algorithms. Furthermore, RNA expression is not fully predictive of protein abundance levels due to differential translational regulation (13), so Visium-IF could be useful for proteins where RNA quantification is not sufficient, such as extracellular matrix proteins or secreted factors.

Several software packages have been developed in the Python and R programming languages for cell segmentation and subsequent registration of microscopy images to anatomical reference atlases (14,15). While some of these frameworks have been applied to spatially resolved transcriptomics data (16), they were designed and optimized for cultured cells and mouse brain tissue sections. In contrast, other packages, such as Fiji (17) and CellProfiler (18) lack functions to automatically split large whole slide images into individual arrays, and have limited features for visualizing large images.

To address these challenges and provide an end-to-end solution that is tailored to analyzing images for the Visium platform, we developed *VistoSeg. VistoSeg* is a freely-available MATLAB-based software package that facilitates segmentation, analysis, and visualization of histological and immunofluorescence images generated on the Visium platform for integration with gene expression data. We also provide user-friendly tutorials and vignettes to support application of *VistoSeg* to new datasets generated by the scientific community.

## Results

Here we describe implementation and requirements for *VistoSeg*, an automated pipeline for processing high-resolution histological or immunofluorescent images acquired during the Visium or Visium-IF workflows for integration with downstream spatial transcriptomics analysis.

### Visium

In the Visium workflow, tissue is cryosectioned and collected into individual arrays of a Visium gene expression slide (**Fig. 1**). For example, we previously collected and analyzed brain tissue sections from the dorsolateral prefrontal cortex (DLPFC) that contained the six cortical layers and white matter (**Fig. 2A**) (6). Here, in a separate DLPFC sample we stained the tissue with H&E and acquired bright field histology images using a slide scanner (**Fig. 2B**). We next describe the capabilities and functions of each section of the *VistoSeg* pipeline.

**Figure 2:**
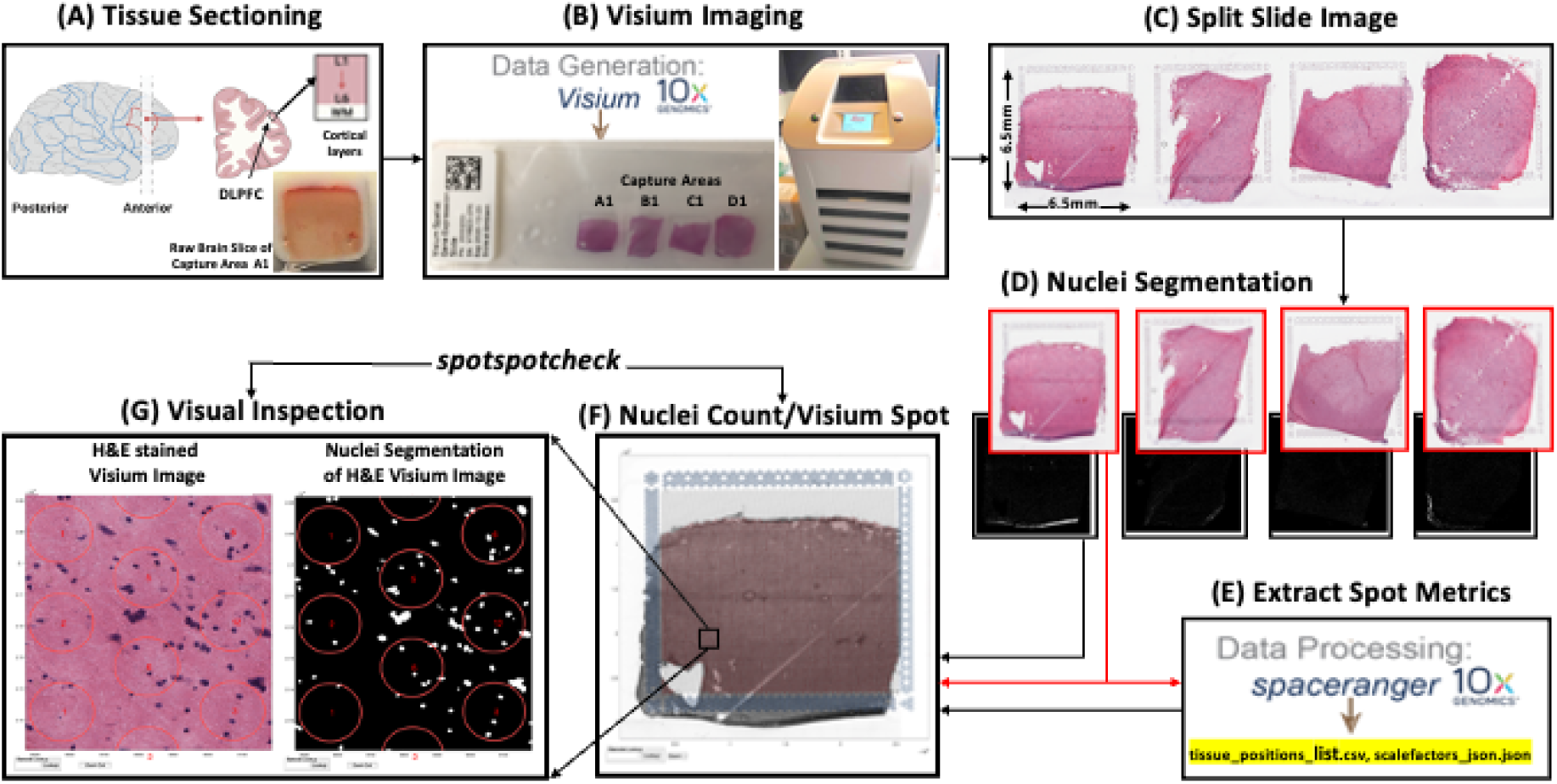
Visium image processing workflow: (**A**) The data presented here is from tissue sections obtained from the human dorsolateral prefrontal cortex (DLPFC). Each tissue section spans the six cortical layers plus the white matter. (**B**) Tissue sections were collected on the ‘Visium gene expression slide’, fixed, stained, and scanned using a slide scanner (Leica CS2). (**C**) The CS2 scanner produced a large high resolution TIFF file of the entire slide, which was split into individual arrays/capture areas using splitSlide(). (**D**) A two-step process using VNS()and refineVNS()was used for segmenting nuclei for each individual capture area. (**E**) Using the split images from (D), Space Ranger exports the spot metrics (Visium spot diameter, spot spacing and spot coordinates) in the ‘tissue_positions_list.csv’ and ‘scalefactors_json.json’ files. (**F**) countNuclei() used the outputs from Space Ranger and computed the nuclei count per Visium spot which were exported in the ‘tissue_spot_counts.csv’ file. (**G**) The *spotspotcheck* GUI allowed for visual inspection of the segmented nuclei and its count per spot by toggling between the segmented binary image and raw histology image. This GUI allows users to zoom in/zoom out on the high resolution image and has a search tab to look for a specific Visium spot of interest based on the spot’s barcode identifier. This utility is useful for exploring image features that might be related to observed gene expression patterns.

### Split Visium slide image

Processing of the Visium data through Space Ranger by 10x Genomics (19) requires individual images for each capture area. The final TIFF images exported from the slide scanner contained the whole slide (**Fig. 2C**), and needed to be split into the individual capture areas to run Space Ranger (19). Obtaining individual capture area images was performed in *VistoSeg* using the splitSlide() function. The function reads in the TIFF image as an RGB (Red, Green, Blue) matrix and splits it along the X-axis into 4 equal RGB matrices. Each individual RGB matrix is considered a capture area and is saved as a separate RGB TIFF file at 70% resolution of the raw individual RGB matrix. For example, a matrix that is of size 10,000 × 10,000 × 3 pixels in size is compressed to 7,000 × 7,000 × 3 pixels. This compression was useful to accommodate file size limitations in MATLAB, which could not store data larger than 2^32^-1 bytes in an image (TIFF, PNG, JPEG), which otherwise can be easily exceeded by the split RGB matrices. We also ensured that the compressed matrices/images were at least 2,000 pixels in either dimension (X and Y) as per Space Ranger’s input recommendations (20).

### Segmentation

Nuclei counts for each Visium spot can be used for downstream gene expression and spot deconvolution analysis. The *Visium Nuclei Segmentation* VNS() function was used to segment nuclei (**Fig. 3B-D**) from the raw histology image (**Fig. 3A**); VNS() first performed gaussian smoothing and contrast adjustment on the raw histology image (**Fig. 3A**) to enhance the nuclei in the image (**Fig. 3B**). The enhanced image was converted from RGB color space to CIELAB color space, also called L*a*b color space (**Fig. 3C**). L*a*b color space is defined by: L, luminosity layer measures lightness from black to white; a, chromaticity layer measures color along the red–green axis; and b, chromaticity layer measures color along blue–yellow axis. The CIELAB color space quantifies the visual differences caused by the different color gradients in the image. The a*b color space was extracted from the L*a*b-converted image and used for *K*-means clustering with *imsegkmeans* (**Fig. 3D**), along with the number of color gradients (*k*) that were visually identified in the image. The function created a binary mask for the (*k*) distinguishable color gradients in the image. Given that nuclei in the histology images had a bright color that could be easily differentiated from the background, a binary mask generated for the nuclei color gradient was used for initial nuclei segmentation (**Fig. 3D** cluster 3, **Fig. 3D`**).

**Figure 3:**
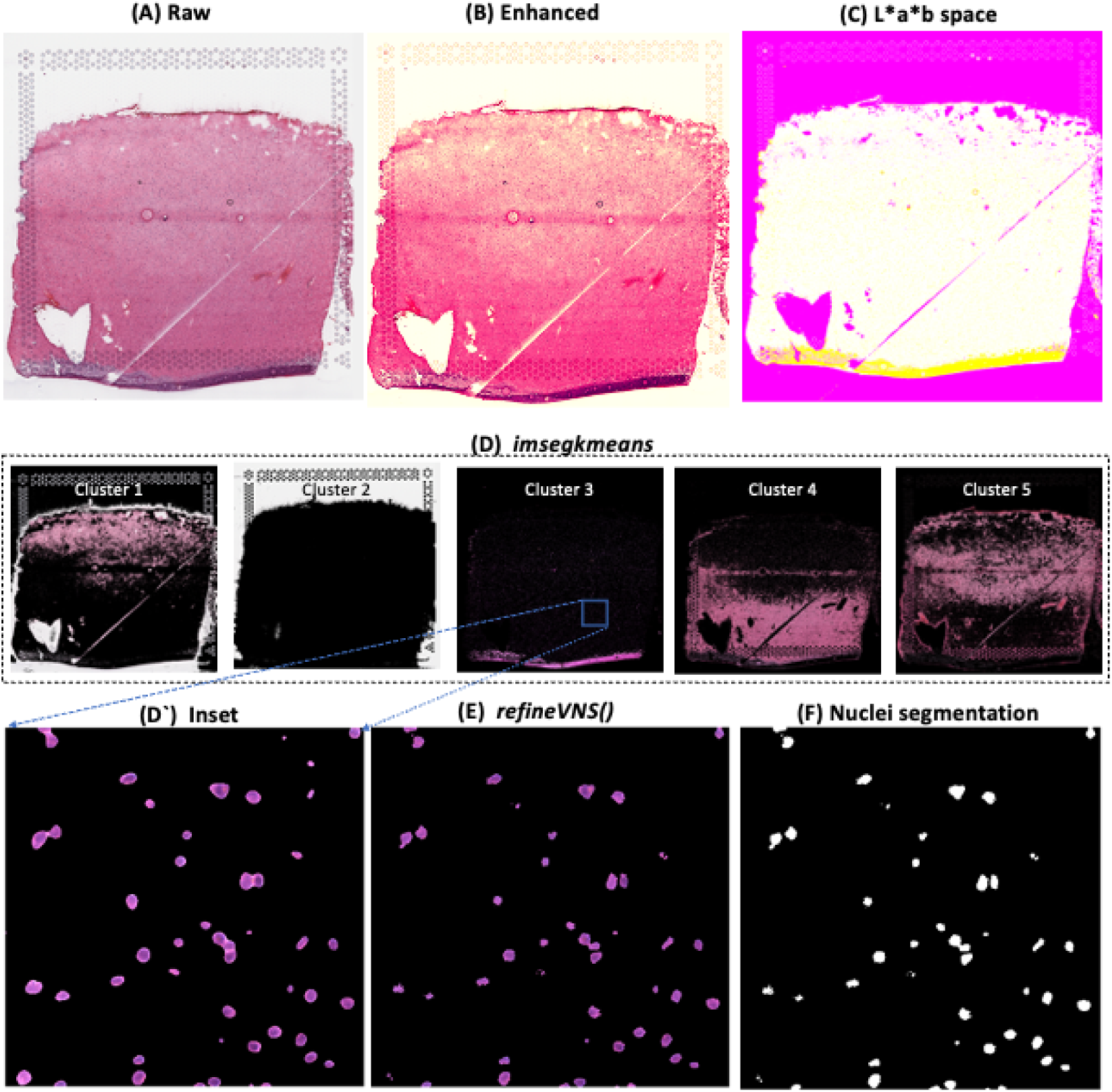
Visium segmentation workflow. (**A**) Raw histology image of human dorsolateral prefrontal cortex (DLPFC). (**B**) Gaussian smoothed and contrast adjusted image of the raw histology image in (A). (**C**) The enhanced image from (B) converted from RGB color space to L*a*b color space. (**D**) Different color gradients (*k*=5) identified by *K*-means clustering function *imsegkmeans* applied to the raw histology image. Cluster 3 corresponded to the nuclei, stained blue in the raw histology image. (**D`**) An inset of nuclei-cluster 3 in (D). (**E**) Output of refineVNS() from nuclei-cluster 3 (D`). (**F**) Final binary nuclei segmentation obtained from (E).

Due to the smoothening performed by the VNS() function, the nuclei edges were blurred and the nuclei that are in close proximity were segmented as a single region (**Fig. 3D`**). To further refine these segmentations for accurate detection of individual nuclei (**Fig. 3F**), we used the refineVNS() function. This function extracted the intensity of the pixels from the binary mask of nuclei generated by VNS(), and applied intensity thresholding (21) to separate the darker nuclei regions at the center from the lighter regions at the borders (**Fig. 3E**).

### spotspotcheck

The *spotspotcheck* Graphic User Interface (GUI, **Fig. 2G**) allows for interactive visual inspection of the nuclei segmentations using the output from Space Ranger software (19). This GUI reconstructs and overlays the spot grid onto the histological image and binary segmentation to display the nuclei count. It allows the user to 1) toggle between the nuclei segmentation and histology images, 2) search for spots with specific spatial barcode IDs, and 3) zoom in and out to a specific location on the image. The user can obtain the nuclei count information either (i) by checking the ‘Get cell counts’ option in the *spotspotcheck* start window or (ii) by using the countNuclei() function from the pipeline. Both options provide a final table, exported as a *CSV* file, with nuclei count per Visium spot for each array, which is used in downstream analyses. *spotspotcheck* enables users to perform bidirectional visual inspection. Thus, the user can evaluate image features including cell densities, morphological features in the raw high resolution histology image, verification of segmentation accuracy across fluorescent channels, identification of imaging or tissue artifacts. Alternatively, the user can visually inspect if gene expression patterns are related to any image features by querying specific spots through their barcode identifier.

### Visium-Immunofluorescence (IF)

The Visium-IF workflow differs from the Visium H&E workflow on the initial tissue staining steps. In the Visium-IF workflow, samples are not stained with H&E, but are instead subjected to immunofluorescence (IF) antibody labeling to detect proteins of interest, thereby generating proteomic data that can be quantified and integrated with transcriptomic data.

We processed 4 individual tissue sections of the human dorsolateral prefrontal cortex (DLPFC) spanning the six cortical layers (L1-L6) and white matter from 4 neurotypical donors (**Fig. 4**). To visualize cellular composition and cell-type distribution across the tissue section, these samples were immunostained with four different cell-type markers: NeuN for neurons, TMEM119 for microglia, GFAP for astrocytes, and OLIG2 for oligodendrocytes. The resulting immunofluorescence images were acquired using a Vectra Polaris slide scanner (Akoya Biosciences) and processed in Phenochart (22) and inForm (23) to decompose the multi-spectral profiles (**Fig. 4A**). The final image outputs were spectrally unmixed multichannel TIFF tiles of the entire slide (∼600 tiles). After linear unmixing, we used InformStitch() to stitch together individual tiles and recreated a multichannel TIFF image spanning the whole slide (**Fig. 4B**). InformStitch() used the x and y coordinates of each tile, as saved in the filename, to perform stitching. Next, splitSlide_IF() splitted (along the Y-axis) the stitched image from InformStitch() into 4 individual capture area images in multichannel TIFF format (**Fig. 4C**). We then used *dotdotdot* (24) to perform segmentations of each fluorescent channel to identify Regions of Interest (ROIs, **Fig. 4D)**. Finally, countNuclei() quantified the size, intensity and location of segmented signals in each channel for integration with gene expression data (**Fig. 4E**). The output table from countNuclei() has two columns per channel by default: (1) count of segmented ROIs per Visium spot and (2) percentage of the spot covered by the segmented signal. Other user-defined metrics, including mean fluorescent intensity per spot or mean intensity of segmented ROIs for each channel within a spot, can also be extracted.

**Figure 4:**
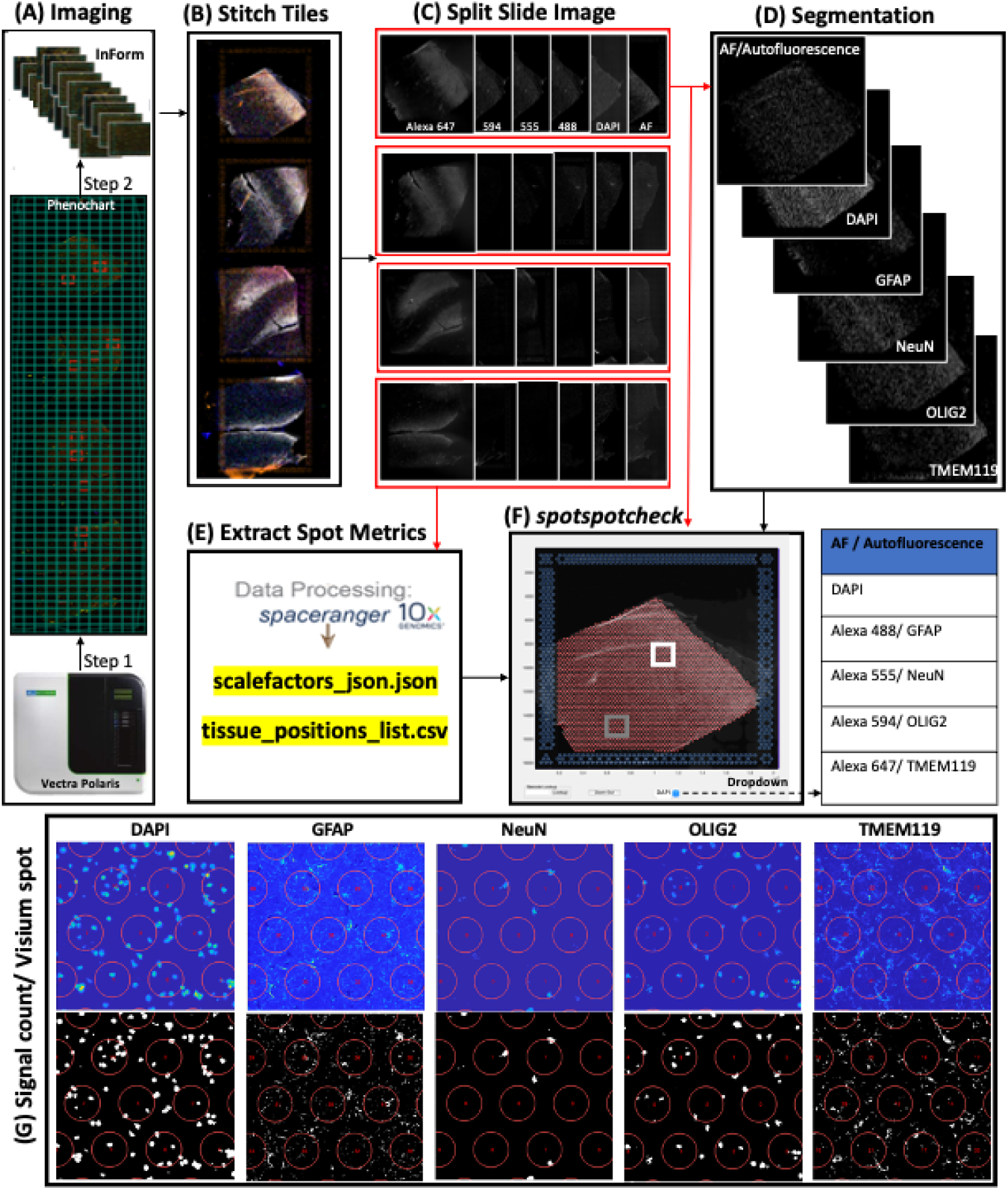
Visium-IF image processing workflow. (**A**) Multispectral immunofluorescent images of the gene expression slide from the Visium-IF workflow were acquired using a Vectra Polaris slide scanner. A single Region Of Interest (ROI) covering all arrays on the slide was selected through Phenochart and this ROI was split into multiple tiles which were spectrally unmixed into multichannel TIFFs using inForm. (**B**) After unmixing, the tiles from (A) were stitched to recreate the multi-channel TIFF of the whole slide. (**C**) The recreated multi-channel TIFF was split into individual arrays using splitSlide_IF(). (**D**) Segmentation of each channel was obtained using *dotdotdot* (24) (example showing capture area A1). (E) Using the split images from (C) Space Ranger extracted spot metrics (Visium spot diameter, spot spacing and spot coordinates) and outputs them in the ‘tissue_positions_list.csv’ and ‘scalefactors_json.json’ files. (**F**) The *spotspotcheck* GUI provides a dropdown menu for different channels in the multichannel TIFF (DAPI, GFAP, NeuN, OLIG2, TMEM119 in this sample). It allows for visual inspection by hovering over different regions in the image. For example, we explored the white matter (white square) and gray matter (gray square) in this DLPFC sample. (**G**) The signal count per Visium spot computed by countNuclei() on the white matter (white square in F). Similar quantification for gray matter is shown in Supplementary Figure 1.

The *spotspotcheck* GUI also supports immunofluorescence images and provides a dropdown menu (**Fig. 4F**,**G**) to select different channels in the multichannel TIFF image along with the other functionalities described in the *spotspotcheck* section. We observed accurate quantification in white matter of the DLPFC (**Fig. 4G**), where we expected higher counts for glial cells stained by TMEM119 (microglia), GFAP (astrocytes) and OLIG2 (oligodendrocytes) compared to the neuron-enriched gray matter (**Sup Fig.1**).

### Example use cases

To demonstrate the utility of *VistoSeg* on datasets obtained from different tissue types, we applied *VistoSeg* to public datasets provided by 10x Genomics (25). First, we ran the *VistoSeg* pipeline on mouse brain tissue sample featuring differing cell densities across the image (**Sup Fig. 2A**). The default settings (smoothing and *k*=5 color clusters in the VNS() function) were used to obtain accurate nuclei segmentations across different regions (inset in green showing cortex and inset in red showing hippocampal pyramidal cell layer, **Sup Fig. 2B**) on the low resolution raw histology image (**Sup Fig. 2A**).

We also applied *VistoSeg* to a human heart tissue sample featuring low cell density and a slightly different background coloration pattern (**Sup Fig. 3**). In this sample, nuclei were surrounded by both the white and pink regions, as opposed to the human brain images where nuclei are almost always surrounded by a pink neuropil region (**Fig. 3**). The smoothening step was not performed to avoid blurring of cell borders as the cells were faintly stained (**Sup Fig. 3A**) and we verified segmentation quality on the *spotspotcheck* GUI (**Sup Fig. 3A-B**).

## Discussion

*VistoSeg* leverages the MATLAB Image Processing Toolbox (26) to provide user-friendly functionality for processing, analysis, and interactive visualization of both histological and fluorescent images generated in conjunction with the 10x Genomics Visium platform. A major feature of the *VistoSeg* pipeline is the quantification of detected signal in the RGB image in the Visium workflow or in each individual fluorescent channel in the Visium-IF workflow. It extracts multiple user-defined metrics including number of cells per spot, percentage spot covered with cells or pathology, mean fluorescence intensity of spot and mean fluorescence intensity of the segmented regions in the spot.

This quantitative output from *VistoSeg* can be integrated with spatial gene expression data to improve spot level resolution and add biological insights to downstream analyses. For example, using histological images of human DLPFC analyzed with *VistoSeg*, we identified “neuropil” spots lacking cell bodies, which we hypothesized were enriched for neuronal processes. We confirmed this hypothesis by demonstrating that these “neuropil” spots are enriched for genes preferentially expressed in the transcriptome of synaptic terminals (6). *VistoSeg* can also be used to identify spots containing a single cell for restricted analysis to individual cell types, or spots containing pathology for analysis related to disease diagnosis. For example, spatial transcriptomics has been used to identify gene expression changes associated with amyloid beta pathology in Alzheimer’s disease (8).

*VistoSeg* was designed to address an image analysis gap in the current Visium processing pipeline available from 10x Genomics. However, we note some limitations such as large memory requirements (∼75 GB) for loading and saving images, and the VNS() function requires manual user inputs and cannot currently be fully automated. However, we similarly note that existing image analysis softwares, such as HALO (3), Squidpy (27) also have limitations on processing times and parametrization, and there are currently no available modules to support integration of gene expression data with segmented images.

While we recognize that MATLAB is closed-source, it is compatible with open science (28) and readily available to academic users. MATLAB supports reading images from various proprietary file formats from multiple instrument manufacturers. All the code and data used by *VistoSeg* is freely available (Data Availability) and the main output from *VistoSeg* is in CSV file format that can be easily incorporated into R objects like SpatialExperiment (29), Seurat (30), or Python objects like AnnData (31), which are two commonly used languages for analysis of spatial transcriptomic data. In addition, conversion utilities like zellkonverter (32) facilitate intercommunication among these programming languages. As other packages are made available that expand on the key infrastructure provided by SpatialExperiment, *VistoSeg* will be compatible with them.

*VistoSeg* output could be further integrated with softwares such as VAMPIRE (33) to identify and classify specific cell types based on nuclear or cellular morphology. We further anticipate that outputs of *VistoSeg* can be used in the future to calculate parameters such as cell density for incorporation into unsupervised clustering approaches. Clustering algorithms have already begun to leverage imaging data (34), and we anticipate that more quantitative outputs from softwares such as *VistoSeg* can improve identification of biologically relevant spatial domains.

## Conclusion

We have developed *VistoSeg* as a comprehensive and user-friendly image processing toolkit, which is optimized for Visium and Visium-IF workflows, to facilitate integration of the rich anatomical and proteomic data found in histological or fluorescent images with corresponding spatial transcriptomic data. *VistoSeg* performs automatic splitting of whole slide images for down stream data processing and allows for visualizing the individual high resolution raw histology and immunofluorescence images. The pipeline is easily adaptable for images obtained from Visium workflows from different tissues, organs and species. The pipeline is available http://research.libd.org/VistoSeg and includes a detailed tutorial with example data for learning how to use *VistoSeg*.

## Materials and Methods

### Tissue Preparation and Image Acquisition for Visium

Hematoxylin and eosin (H&E) staining was performed on fresh frozen tissue according to manufacturer’s instructions to identify nuclei (dark blue/purple) and cytoplasm (pink) in the tissue section. The two stains combine to label features of the tissue in various shades of pink and blue. Thus, the number of colors present will depend on the cellular composition of the tissue. Following H&E staining, the Visium slide was imaged on a Leica Aperio slide scanner (**Fig. 1B**) equipped with a color camera and a 20X/0.75NA objective with a 2X optical magnification changer, which meets the recommended microscopy specification outlined by Visium Spatial Gene Expression Imaging Guidelines from 10x Genomics (35). This protocol produced high-resolution (0.253 µm per pixel) images for downstream analysis.

### Immunofluorescence Staining and Image Acquisition for Visium-IF

Immunofluorescence (IF) staining was performed according to the manufacturer’s instruction (catalog no.CG000312 Rev C, 10x Genomics). Briefly, the human dorsolateral prefrontal cortex (DLPFC, n=4) was microdissected and cryosectioned at 10-micron thickness. Sections were mounted on a Visium Spatial Gene Expression Slide (catalog no. 2000233, 10x Genomics), fixed in pre-chilled methanol, blocked in BSA-containing buffer, and incubated for 30 minutes at room temperature with primary antibodies against NeuN, TMEM119, GFAP, and OLIG2 (mouse anti-NeuN antibody conjugated to Alexa 488 (Sigma Aldrich, Cat# MAB377X, 1:100), rabbit anti-TMEM119 antibody (Sigma Aldrich, Cat# HPA051870, 1:20), rat anti-GFAP antibody (Thermofisher, Cat# 13-0300, 1:100), and goat anti-OLIG2 antibody (R&D systems, Cat# AF2418, 1:20). Following washes, appropriate secondary antibodies were applied for 30 minutes at room temperature (donkey anti-rabbit IgG conjugated to Alexa 555 (Thermofisher, Cat# A-31572, 1:300), donkey anti-rat IgG conjugated to Alexa 594 (Thermofisher, Cat# A-21209, 1: 600), and donkey anti-goat IgG conjugated to Alexa 647 (Thermofisher, Cat# A-21447, 1:400)). DAPI (Thermofisher, Cat# D1306, 1:3000, 1.67 μg/ml) was applied for nuclear counterstaining. The slide was coverslipped with 85% glycerol and imaged on a Vectra Polaris slide scanner (Akoya Biosciences) at 20x magnification with the following exposure time per given channel: 2.1 msec for DAPI; 143 msec for Opal 520; 330 msec for Opal 570; 200 msec for Opal 620; 1070 msec for Opal 690; 100 msec for Autofluorescence (AF) prior to downstream transcriptomics.

### VistoSeg

*VistoSeg* has been tested on Linux, Windows and MacOS. MATLAB 64-bit version R2019a or later with the Image Processing Toolbox (26) is required to run the *VistoSeg*. The multiplane, high-resolution whole slide images that we have typically acquired on different brain regions using Visium and Visium-IF workflows, are ∼25GB (TIFF format). The system RAM should approximately be thrice the size of the input raw image to load it into MATLAB (∼75GB), prior to splitting the image into the individual capture areas. The remainder of the processing that is carried out on the individual capture TIFFs can be performed on a system with 16GB of RAM. The pipeline is available at https://github.com/LieberInstitute/VistoSeg (36). Steps to run the pipeline and sample images are provided in the *VistoSeg* website http://research.libd.org/VistoSeg/.

### Space Ranger

The 10x Genomics Space Ranger software (19) uses the output TIFF files produced by the splitSlide() and splitSlide_IF() functions as input for quantification of gene expression in a spatial context (**Fig. 2E, 4E**). This software generates a *CSV* and a *JSON* file that contains metrics to reconstruct the spot grid in *spotspotcheck*. The different inputs required and the workflow of Space Ranger (19) are explained in detail in the following website chapter http://research.libd.org/VistoSeg/step-3-spaceranger.html.

## Acknowledgements

We thank Anthony Ramnauth (LIBD) and Uma Kaipa (LIBD) for testing the code functionality. We thank the “spatialLIBD” team (LIBD and JHU) for their feedback on *VistoSeg* and testing it across multiple datasets. Finally, we thank the families of the decedents, who donated the brain tissue used in this study as well as donors for the public datasets we used in this manuscript. A preprint of this work is available on bioRxiv: doi: https://doi.org/10.1101/2021.08.04.452489.

## Data Availability

Examples of code, data, output and results are available at http://research.libd.org/VistoSeg/index.html#data-availability https://github.com/LieberInstitute/VistoSeg (36). All inputs and outputs are available through Figshare (37). Public datasets provided by 10x Genomics (25).

## Supplementary Material

## Supplementary Figures

**Supplementary Figure 1:**
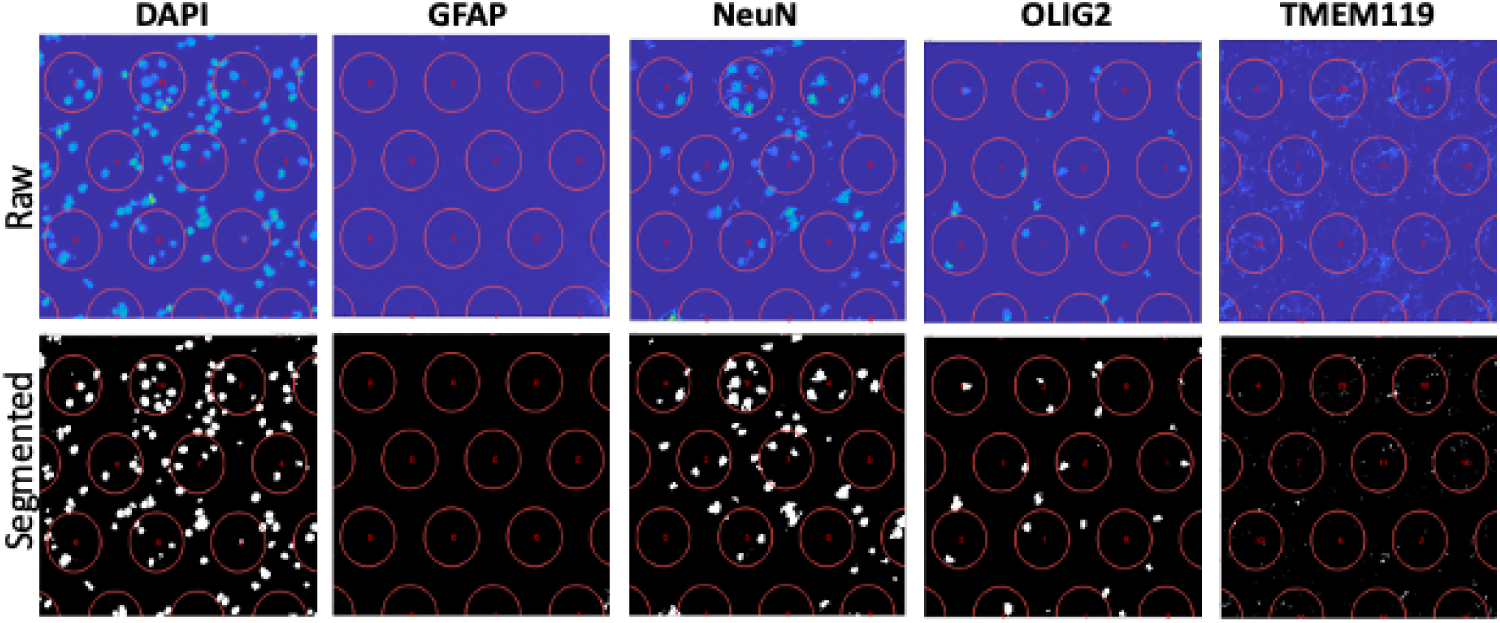
*spotspotcheck* on gray matter. *spotspotcheck* visualization and quantification (DAPI, GFAP, NeuN, OLIG2, TMEM119) from a representative field of gray matter from the DLPFC sample shown in **Figure 4**. Top row shows the insets of raw Visium-IF images and the bottom row shows the respective fluorescent segmentations obtained from *dotdotdot* (24).

**Supplementary Figure 2:**
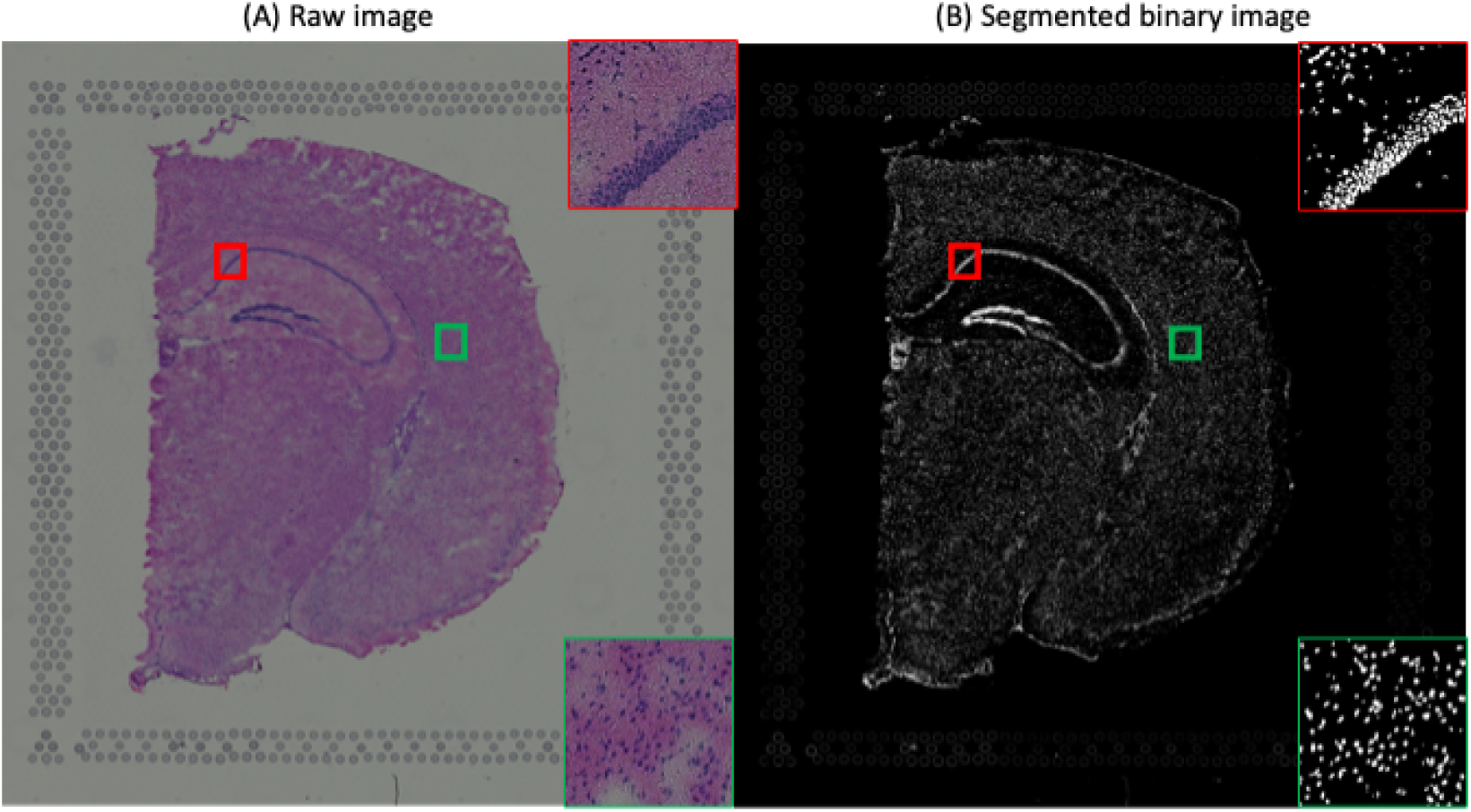
Mouse brain coronal section. (**A**) Raw hematoxylin and eosin (H&E) image of a fresh frozen mouse brain tissue section in the coronal plane. Insets showing cell densities at different regions in the section. (**B**) Final segmented binary image of (A) from the refineVNS()function. Red and green squares are random insets that are shown at higher resolution.

**Supplementary Figure 3:**
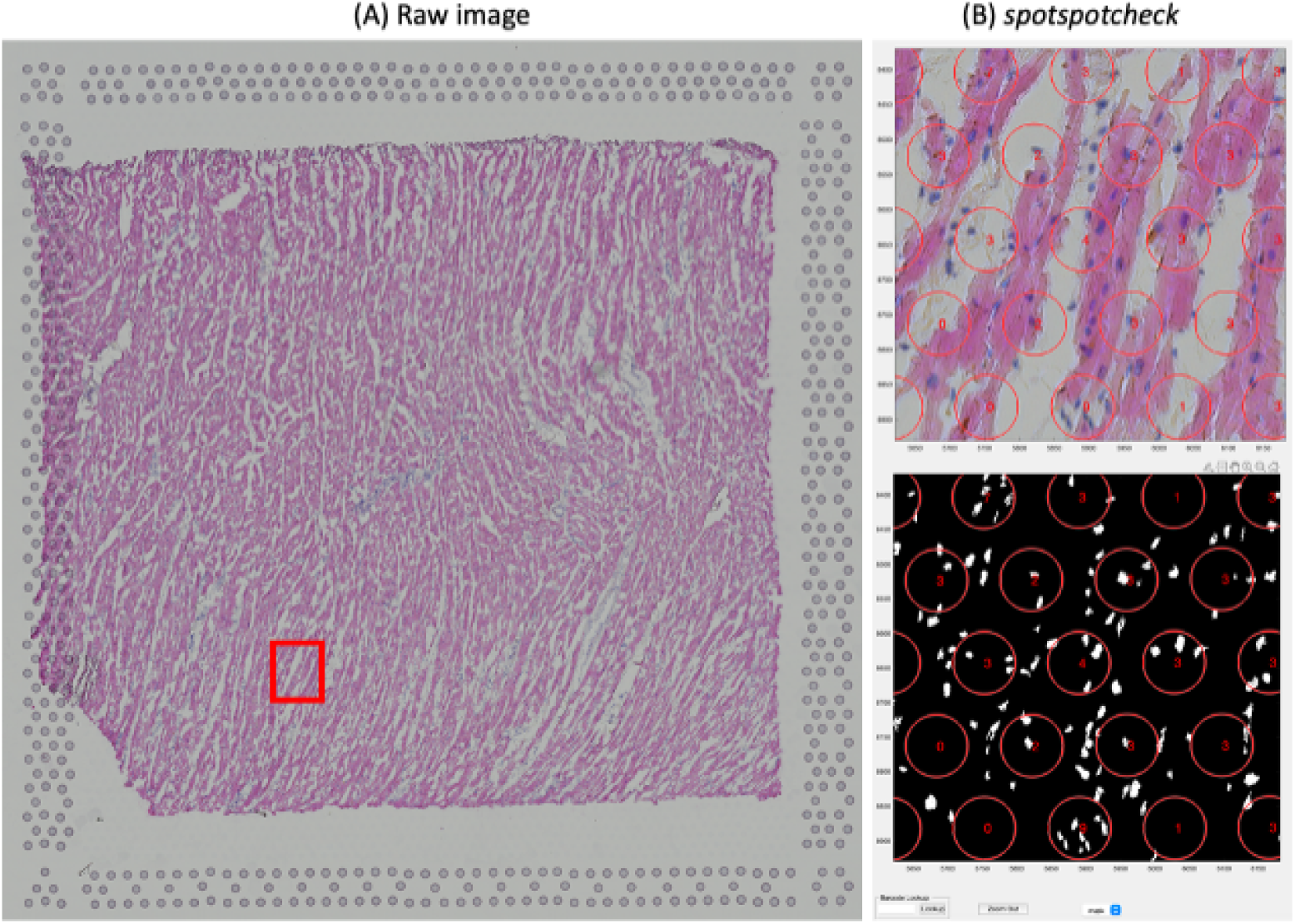
Human heart tissue. (**A**) Raw hematoxylin and eosin (H&E) image of a fresh frozen human heart tissue section. (**B**) Snapshot of *spotspotcheck* visualization of a random region in (A) showing segmentation and quantification.

## Supplementary Files

These files are available through Figshare at https://doi.org/10.6084/m9.figshare.19729567.v1 (37).

**Supplementary File 1: Human_DLPFC_IF.zip**. This folder contains files to run *spotspotcheck* on the human DLPFC IF sample. These files are:

- **scalefactors_json.json**. JSON file from Space Ranger providing the spot diameter estimation in pixels for the high resolution raw histology image. See https://support.10xgenomics.com/spatial-gene-expression/software/pipelines/latest/using/count for more information about this file.
- **tissue_positions_list.csv**. CSV file containing spot barcode, if the spot was called under (1) or out (0) of tissue, the array position, image pixel position x, and image pixel position y for the high resolution raw histology image. See https://support.10xgenomics.com/spatial-gene-expression/software/pipelines/latest/using/count for more information about this file.
- **tissue_spot_counts.csv**. CSV file containing all the information in tissue_positions_list.csv and also the signal count and percentage area of signal per spot.
- **V10B01-087_A1_segmentation.mat**. MATLAB mat file consisting segmentations of all channels (AF-Autofluorescence, DAPI, GFAP, NeuN, OLIG2, TMEM119) from the raw TIFF image of human DLPFC stored as V10B01-087_A1.tif
- **V10B01-087_A1.tif**. Raw multichannel (AF-Autofluorescence, DAPI, GFAP, NeuN, OLIG2, TMEM119) TIFF image of human DLPFC sample used in Figure 4.

**Supplementary File 2: Mouse_brain.zip**. This folder contains files to run *spotspotcheck* on the mouse brain H&E sample. These files are:

- **scalefactors_json.json**. JSON file from Space Ranger providing the spot diameter estimation in pixels for the high resolution raw histology image. See https://support.10xgenomics.com/spatial-gene-expression/software/pipelines/latest/using/count for more information about this file.
- **tissue_positions_list.csv**. CSV file containing spot barcode, if the spot was called under (1) or out (0) of tissue, the array position, image pixel position x, and image pixel position y for the high resolution raw histology image. See https://support.10xgenomics.com/spatial-gene-expression/software/pipelines/latest/using/count for more information about this file.
- **V1_Adult_Mouse_Brain_image.tif**. Raw H&E histology image of mouse brain tissue section.
- **V1_Adult_Mouse_Brain_image_cluster.mat**. MATLAB mat file with different color gradient clusters obtained from V1_Adult_Mouse_Brain_image.tif image.
- **V1_Adult_Mouse_Brain_image_nuclei.mat**. MATLAB mat file with nuclei segmentation from V1_Adult_Mouse_Brain_image.tif.

**Supplementary File 3: Human_heart.zip**. This folder contains files to run *spotspotcheck* on the human heart H&E sample. These files are:

- **scalefactors_json.json**. JSON file from Space Ranger providing the spot diameter estimation in pixels for the high resolution raw histology image. See https://support.10xgenomics.com/spatial-gene-expression/software/pipelines/latest/using/count for more information about this file.
- **tissue_positions_list.csv**. CSV file containing spot barcode, if the spot was called under (1) or out (0) of tissue, the array position, image pixel position x, and image pixel position y for the high resolution raw histology image. See https://support.10xgenomics.com/spatial-gene-expression/software/pipelines/latest/using/count for more information about this file.
- **tissue_spot_counts.csv**. CSV file containing all the information in tissue_positions_list.csv and also the signal count and percentage area of signal per spot.
- **V1_Human_Heart_image.tif**. Raw H&E histology image of human heart tissue section.
- **V1_Human_Heart_image_cluster.mat**. MATLAB mat file with different color gradient clusters obtained from V1_Human_Heart_image.tif.
- **V1_Human_Heart_image_nuclei.mat**. MATLAB mat file with nuclei segmentation from V1_Human_Heart_image.tif

